# A scalable sparse neural network framework for rare cell type annotation of single-cell transcriptome data

**DOI:** 10.1101/2022.06.22.497193

**Authors:** Yuqi Cheng, Xingyu Fan, Jianing Zhang, Yu Li

## Abstract

Cell type annotation is critical to understand the cell population heterogeneity in the single-cell RNA sequencing (scRNA-seq) analysis. Due to their fast, precise, and user-friendly advantages, automatic annotation methods are gradually replacing traditional unsupervised clustering approaches in cell type identification practice. However, current supervised annotation tools are easily overfitting, thus favoring large cell populations but failing to learn the information of smaller populations. This drawback will significantly mislead biological analysis, especially when the rare cell types are important. Here, we present scBalance, an integrated sparse neural network framework that leverages the adaptive weight sampling and dropout techniques for the auto-annotation task. Using 20 scRNA-seq datasets with different scales and different imbalance degrees, we systematically validate the strong performance of scBalance for both intra-dataset and inter-dataset annotation tasks. Furthermore, we also demonstrate the scalability of scBalance on identifying rare cell types in million-level datasets by uncovering the immune landscape in bronchoalveolar cells. Up to now, scBalance is the first and only auto-annotation tool that expands scalability to 1.5 million cells dataset. In addition, scBalance also shows a fast and stable speed outperforming commonly used tools across all scales of datasets. We implemented scBalance in a user-friendly manner that can easily interact with Scanpy, which makes scBalance a superior tool in the increasingly important Python-based platform.

## Introduction

Since the first establishment of single-cell RNA sequencing (scRNA-seq) by *Tang et al*. in 2009^1^, this technology has rapidly become popular for scientists in various biological research fields. Compared with traditional bulk RNA sequencing, which only measures the average gene expression level of the samples, scRNA-seq provides a powerful method to profile transcriptome on the cell-specific level, enabling analyzing individual cells and giving a more informative insight into cell heterogeneity. The rapid development of scRNA-seq technology has been widely used in several biological research areas, such as cancer research^2, 3^, COVID analysis^4, 5^, developmental biology research^6^, *etc*. In particular, uncovering and identifying cellular populations is one of the critical tasks in single-cell analysis.

Typically, cell type annotation involves two steps: (1) clustering the cells into different subgroups and (2) labeling each group a specific type manually based on the prior-known marker genes. A number of unsupervised machine learning algorithms have been developed, including classical machine learning based methods such as Seurat^7^ and Scanpy^8^, and newly published deep learning based methods, such as scDHA^9^ and CLEAR^10^. However, these methods can be time-consuming and burdensome. For those who do not have too much knowledge of the marker genes, this approach could cost far more time than expected. Automatic cell type annotation methods, in contrast, do not suffer from the manual labeling process. Different from the unsupervised methods, automatic cell-type identification tools are mainly designed on supervised learning frameworks. Taking advantage of this, they are becoming a predominant tool to identify cell types in single-cell experiments. With the unprecedented boom in the well-annotated scRNA-seq atlas and the rapid promotion of the Human Cell Atlas project^11, 12^, auto-annotation tools are facing a more broad prospect than anytime before. Hitherto, thirty-two auto-annotation tools are developed and published^13^. For example, SingleCellNet^14^ utilizes a random-forest classifier to solve the cross-platform and cross-species annotation tasks. And ACTINN^15^ implements a simple artificial neural network to overcome the batch effect.

Although so many tools have been established in recent years, most of those often fail to identify the entire population because of the existence of rare cell types. From the perspective of the cell composition, scRNA-seq datasets are always imbalanced, which have common cell types and rare cell types. The rare population is a small proportion of cells in the single-cell dataset. For example, the dendritic cell usually takes 1% to 5% of peripheral blood mononuclear cells (PBMCs), especially in large datasets^16, 17^. When we train an auto-annotation tool, the classifier is consistently unable to learn their information and thus hard to identify these cell types in the query dataset. However, these rare populations can be crucial, especially in disease research^18^. Meanwhile, we also find that the existing methods have two other main deficiencies. (1) Lack of scalability. Recent scRNA-seq experimental platforms enable investigations of million-level cells^19, 20^. Notably, one of the most recent COVID PBMC atlas has reached 1.5 million cells^17^. Thus computation speed restriction will render auto-annotation packages poorly scalable for the million-level dataset. Moreover, large scale reference datasets add more challenges for learning rare cell types in classifier training, which leads current software more difficult to identify minor groups. Most recently published paper has elevated training scale to 600K cells^21^, however, no published tools successfully report scalability on the million-level cell atlas. (2) Compatibility of the existing tools is not as good as expected. Among the existing Python-based tools, most of the tools such as ACTINN^15^, scPretrain^22^, scCapNet^23^, MarkerCount^24^ are script-based. Considering that Seurat and Scanpy are both packages that can be downloaded from a standard software repository (*e*.*g*., PyPI), running an external Python script on the server will add an additional burden to the user. In addition, some of the tools are no longer maintained or not able to use. All these challenges together make a new annotation tool, which has a balanced ability to label major and minor cell types in a scalable manner, become necessary.

Here, we introduce scBalance, a sparse neural network framework to automatically label rare cell types in all scale scRNA-seq datasets. scBalance leverages the combination of weight sampling and sparse neural network, whereby minor (rare) cell types are more informative without harming the annotation efficiency of the common (major) cell populations. To demonstrate the performance of identifying the rare cell population of our method, we applied scBalance on various real datasets with different imbalance degrees and cell number scales. We also tested the utility of the scBalance on cross-platform tasks to evaluate its ability to learn rare cell type information under batch effect. For comparison, we tested several popular published tools including Scmap-cell^25^, Scmap-cluster^25^, SingleCellNet^14^, SingleR^26^, scVI^27^. Each method represents a traditional machine learning algorithm such as Scmap-cell is based on KNN and SingleCellNet is based on Random Forest. Among them, scBalance remained the most accurate rare populations annotation tool while maintaining high accuracy in annotating major cell types. Across all sizes of the datasets, scBalance also showed fast and stable computation speeds outperforming other approaches. Moreover, scBalance was successfully trained on a published COVID immune cell atlas^17^ (1.5 million cells) and further annotated and discovered new cell types in the published bronchoalveolar lavage fluid (BALF) scRNA-seq dataset^28^. Satisfyingly, our method identified more rare cell types than the original analysis. scBalance is specifically designed to be compatible with Scanpy and Anndata, thus providing a user-friendly application that can be easily downloaded from PyPI and be used as an external API of Scanpy (https://github.com/yuqcheng/scBalance).

## Results

### Overview of the architecture of scBalance

scBalance provides an integrative deep learning framework to perform accurate and fast cell-type annotation, especially on rare cell types, in a scalable manner. (Fig. 1). The structure of the scBalance includes two parts. First, different from all existing tools, we use a specially designed weight sampling technique to adaptively process the imbalanced scRNA-seq dataset. To keep as much information as possible and avoid a huge training time cost, scBalance randomly over-samples the rare populations (minority classes) as well as under-samples the common cell types (majority classes) in each training batch (Fig. 1a). The sampling process is with replacement, and the sampling ratio is adaptive for different reference datasets, which is defined as the cell type proportions in true labels provided by the reference set. By this, we minimize overfitting in the oversampling thus keeping a promising performance of the generalization ability of the scBalance. Meanwhile, regarding the enormous overlapping expression information in the common populations, the under-sampling of the major class enables scBalance to use a relatively small training size with an abundance of training information. Leveraging this design, scBalance yields an exceptional performance in learning features of rare cell types as well as maintains a strong ability in classifying the major cell type, thus also improving its overall annotation accuracy. Moreover, we notice that the reference dataset and the prediction dataset can be generated by different sequencing platforms and protocols such as the 10X platform and Smart-seq platform, thus will naturally introduce a batch effect. In the scBalance, we consider the batch effect a special type of overfitting event and use the dropout^29^ technique to solve this problem. The dropout method we used in the scBalance presents a strong ability to overcome batch effect laying between the reference dataset and the prediction dataset, which increases overall accuracy in cross-platform annotation tasks and decreases time cost compared with other tools. Notably, the dropout layer, due to its excellent capacity of reducing overfitting, also enhances the learning ability of the scBalance to the resampled minor cell types. Companied with the weighted sampling approach, the dropout layer offsets the potentially introduced overfitting in the sampling process thus giving the scBalance a better performance on rare populations identification as well.

**Fig. 1.**
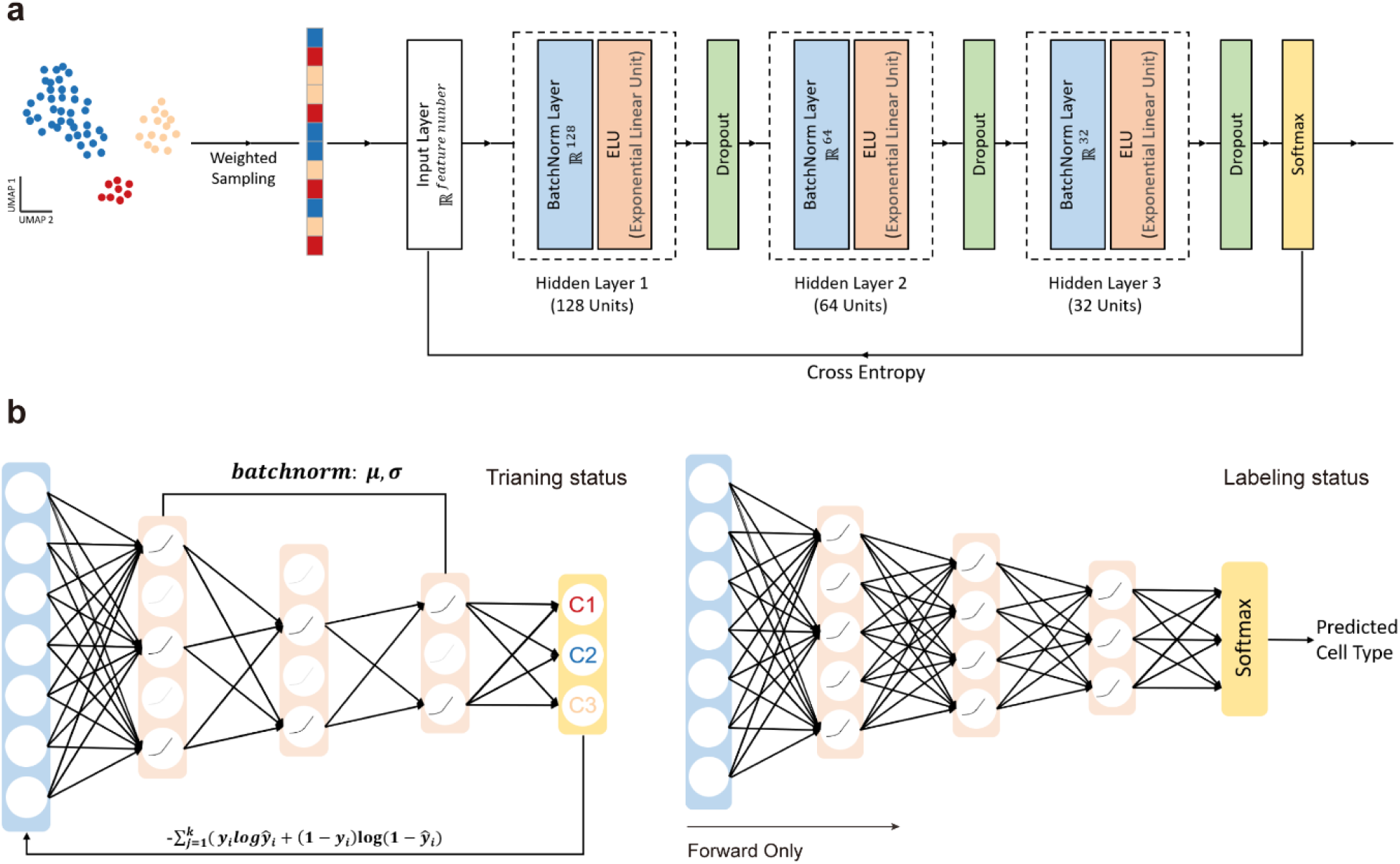
Schematic overview of scBalance. **a**. Structure of scBalance. The method is construct based on supervised learning framework, in which containing a weighted sampling module and a dropout neural network module. Before training, scBalance will automatically choose weight for each cell type in the reference dataset and construct train batch. All genes will be inputted. While training, scBalance will iteratively learn mini batch from three-layer neural network until the cross-entropy loss converges. The dropout layer and sampling technique together improve the rare cell type identification ability by balancing training samples and minimizing overfitting. Batch normalization layer accelerate training speed. **b**. Dropout setting in different stages. In the training stage, scBalance randomly disables neuron in the network. The drop probability is binary with dropping rate=0.5. All the dropped units will be reconnected in the testing stage. The prediction will be processed by a fully connected neural network.

Taken together, scBalance provides a three hidden layers network structure with a batchnorm and dropout setting in each layer. The activation function is set as an exponential linear unit (ELU)^30^ and the output layer as Softmax. In the training mode (Fig. 1b), units in the hidden layer will be randomly disabled so that the neural network will perform like an ensemble classifier; this design can essentially decrease the batch effect to be learned in the training process. In the predicting mode, the network will be set as a fully connected status to keep all parameters being used in the forward process. The model evaluation and backpropagation are based on the cross-entropy loss function and Adam optimizer. To speed up training and predicting process, scBalance also includes a graphics processing unit (GPU) mode besides the default central processing unit (CPU) computation mode. For users with GPU on their server, they can set scBalance to the GPU mode and the running time of the classifier can be reduced by 25% to 30%, with the classification accuracy remains the same.

### scBalance accurately identify rare cell population in the intra-dataset labeling task

We firstly demonstrate the rare cell type identification ability of scBalance in the baseline test. To assess the performance, twelve scRNA-seq datasets with different imbalance degrees and different cell numbers are divided into train set and test set. To make tests more comprehensive, most of the datasets are generated from different sequencing platforms (see Method and Table 1). The true label information of these datasets is only available in evaluating prediction results. Here, we compared scBalance with five methods that are widely used for scRNA-seq cell type identification: SingleCellNet^14^, SingleR^26^, scVI^27^, scmap-cell^25^, and scmap-cluster^25^. To ensure our benchmark comparison is under a fair experiment, we use the uniform preprocessing process for every tool and set all parameters as default. As the true label can be obtained from these 12 datasets, we can quantitatively evaluate the performance of the scBalance and the other five methods by using Cohen’s kappa score (Fig.2a and Supplementary Table 1). According to the result, scBalance outperforms all other methods on all these 12 datasets by having the highest Cohen’s kappa score. Because Cohen’s kappa score provides a minority class sensitive metric, outperforming on this score gives preliminary evidence that the scBalance has more advantages in rare population detecting. Meanwhile, scBalance also shows a stable performance across small-scale datasets (∼1k cells) to mid-scale datasets (∼68k cells) and is robust to different protocols. In particular, on average, scBalance improves over the second-best method, SingleCellNet, by 12.88% regarding the Cohen’s kappa score.

**Table 1.** Description of the 20 datasets used in the experiments. The Dataset column presents the dataset name we used in the article. The cell number column shows the total number of cells in the dataset before preprocessing. The protocol colume shows the sequencing method that generates this dataset. The first 12 datasets are used in the intra-dataset annotation experiment. The following 6 datasets (PBMCBench) are used in the inter-platform annotation experiment and the last two large datasets are used in the scalability experiment.

| Dataset | Cell Number | Class Number | Protocol | Reference |
| --- | --- | --- | --- | --- |
| Deng | 268 | 6 | Smart-seq2 | *Deng et al.*<sup>39</sup> |
| Darmanis | 466 | 9 | SMARTer | *Darmanis et al.*<sup>40</sup> |
| Usoskin | 622 | 4 | STRT-Seq | *Usoskin et al.*<sup>41</sup> |
| CampLiver | 777 | 7 | SMARTer | *Camp et al.*<sup>42</sup> |
| Baron Mouse | 1886 | 13 | inDrop | *Baron et al.*<sup>43</sup> |
| Muraro | 2122 | 9 | CEL-Seq2 | *Muraro et al.*<sup>44</sup> |
| Lake | 3042 | 16 | Fluidigm C1 | *Lake et al.*<sup>45</sup> |
| Baron Human | 8569 | 14 | inDrop | *Baron et al.*<sup>43</sup> |
| Campbell | 21086 | 21 | Drop-Seq | *Campbell et al.*<sup>46</sup> |
| Zilionis | 34558 | 9 | inDrop | *Zilionis et al.*<sup>47</sup> |
| TM<br>(Tabula Muris) | 54865 | 55 | 10X Genomics | *Schaum et al.*<sup>48</sup> |
| Zheng 68K | 65943 | 11 | 10X Genomics | *Zheng et al.*<sup>49</sup> |
| PbmcBench<br>10X (V2) | 6444 | 9 | 10X Genomics<br>(v2) | *Ding et al.*<sup>50</sup> |
| PbmcBench<br>10X (V3) | 3222 | 8 | 10X Genomics<br>(v3) | *Ding et al.*<sup>50</sup> |
| PbmcBench<br>(CEL-Seq) | 253 | 7 | CEL-Seq2 | *Ding et al.*<sup>50</sup> |
| PbmcBench<br>(Drop-Seq) | 3222 | 9 | Drop-Seq | *Ding et al.*<sup>50</sup> |
| PbmcBench<br>(inDrop) | 3222 | 7 | inDrop | *Ding et al.*<sup>50</sup> |
| PbmcBench<br>(Seq-Well) | 3176 | 7 | Seq-Well | *Ding et al.*<sup>50</sup> |
| Cardiac Atlas | 487,106 | 11 | 10X Genomics | *Litviňuková et al.*<sup>31</sup> |
| PKU_Covid Atlas | 1,462,702 | 64 | 10X Genomics | *Ren et al.*<sup>17</sup> |

**Fig. 2.**
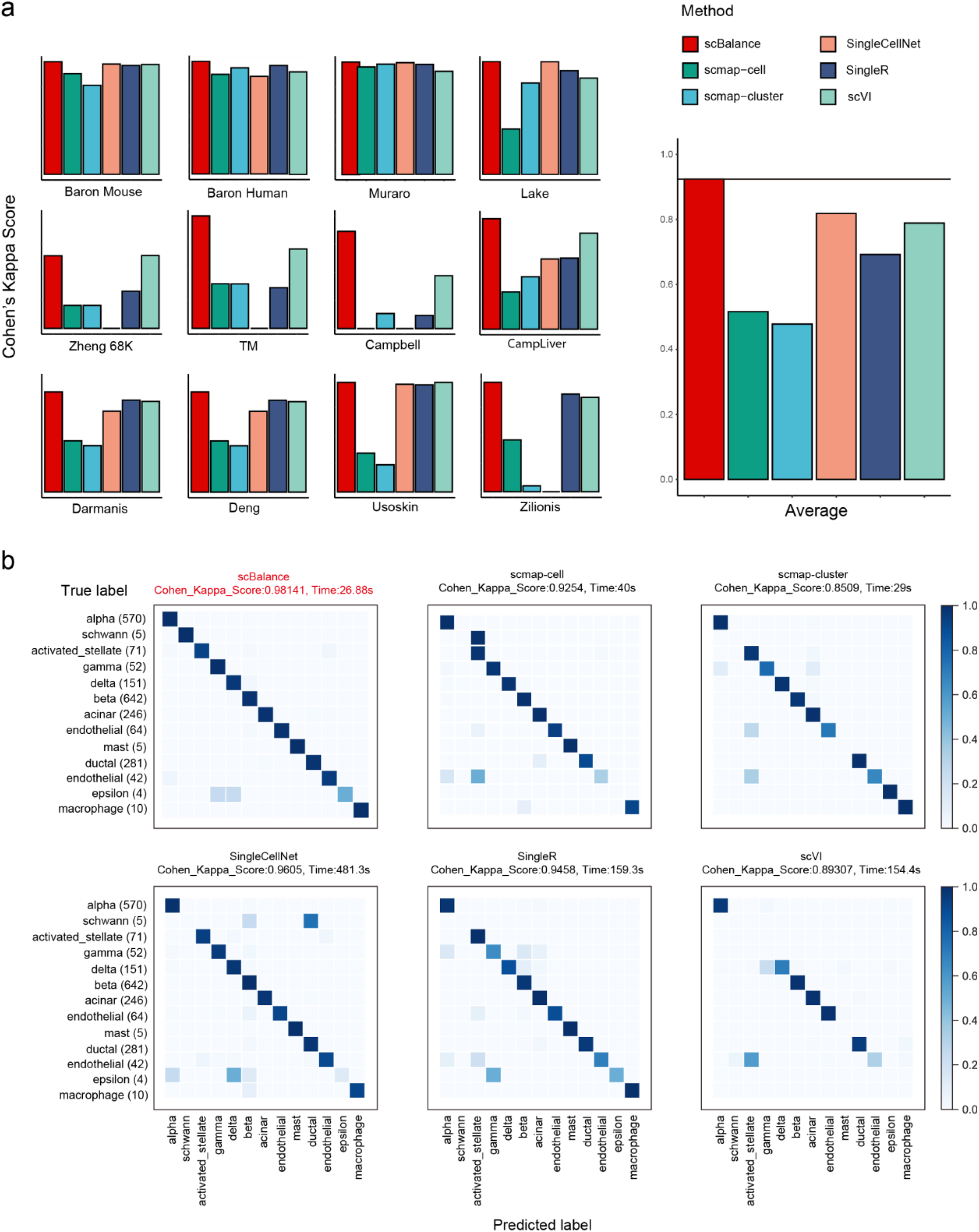
scBalance outperforms existing methods on the rare cell population identification. a. Overall annotation performances (Cohen’s Kappa score) comparing with different methods on various datasets. scBalance shows a stable classification ability on the minor-class sensitive metirc across twelves datasets. b. Confusion matrix of the annotation results on Baron-Human dataset. Only scBalance successfully identifies all the rare type in the testing set. All other methods fail to identify schwann cell. scVI is not able to correctly label all rare cell types.

To better understand how scBalance performs in identifying rare cell types, we generate confusion matrixes on the result of the highly imbalanced datasets for each method (Fig. 2b). As shown in the figures, scBalance successfully identifies both the major and the minor types while the other 5 methods give a poor performance on the minor cell type Significantly, on the Baron Human dataset, which has 13 cell types and the major cell type has 100 times than the minor cell type, scBalance precisely annotates the minor types, including “schwann” cell, “mast” cell, and “epsilon” cell, which only have 5, 5, 4 cells respectively. However, the other five tools fail to identify the part or all of these minor types, even though they all have high accuracy. SingleCellNet gives the most promising Cohen kappa’s score among these five methods, however, it wrongly identifies all the “schwann” cell and the “epsilon” cell. Scmap-cell and scmap-cluster show similar classification results with SingleCellNet. scVI even fails to identify the types with more cells than those 3 minor types but only can recognize the major types with very high cell numbers. Confusion matrix analysis gives primary evidence to evaluate rare cell type classification capability. All these confusion matrixes together indicate that scBalance is a perfect method to identify minor cell types on a scRNA-seq dataset with a high imbalance degree. In summary, scBalance performs well on the baseline annotation task for its stable ability to not only successfully identify the major cell types but also the minor cell types.

### scBalance outperforms in rare population identification under the batch effect

In the realistic scenario, we anticipate that users may train an annotation tool with a dataset generated from a different protocol to that used for the query scRNA-seq profile. However, different sequencing platforms would naturally introduce batch effect into the scRNA-seq dataset, thus adding bias to the data and making rare type annotation much more difficult. To assess the robustness of scBalance to overcome the batch effect and the ability to identify rare cell types on the cross-platform annotation, we used PBMCbench datasets (see Method and Table 1) to test and evaluate the performance. The PBMCbench datasets consist of PBMCs scRNA-seq data generated from different protocols. Given that the 10X chromium and SMART-Seq2 protocols are the most widely used sequencing methods, we mainly use the datasets from these two platforms as the training set and test on the datasets coming from all the other platforms. We still use Cohen’s kappa score evaluate the performance. The results are summarized in the Fig. 3 and Supplementary Table 2. Overall, scBalance achieves the highest scores in all 12 experiments (Fig. 3a). Compared with the second-best method, scBalance elevates the average score from 0.85 to 0.95. The improvement is more significant when the models are trained with SMART-Seq2 dataset, in which scBalance outperforms than SingleCellNet by 13% and scmap-cluster by 56%. Taken together, the results suggest that scBalance has more advantages on the cross-platform annotation tasks than the existing approaches.

**Fig. 3.**
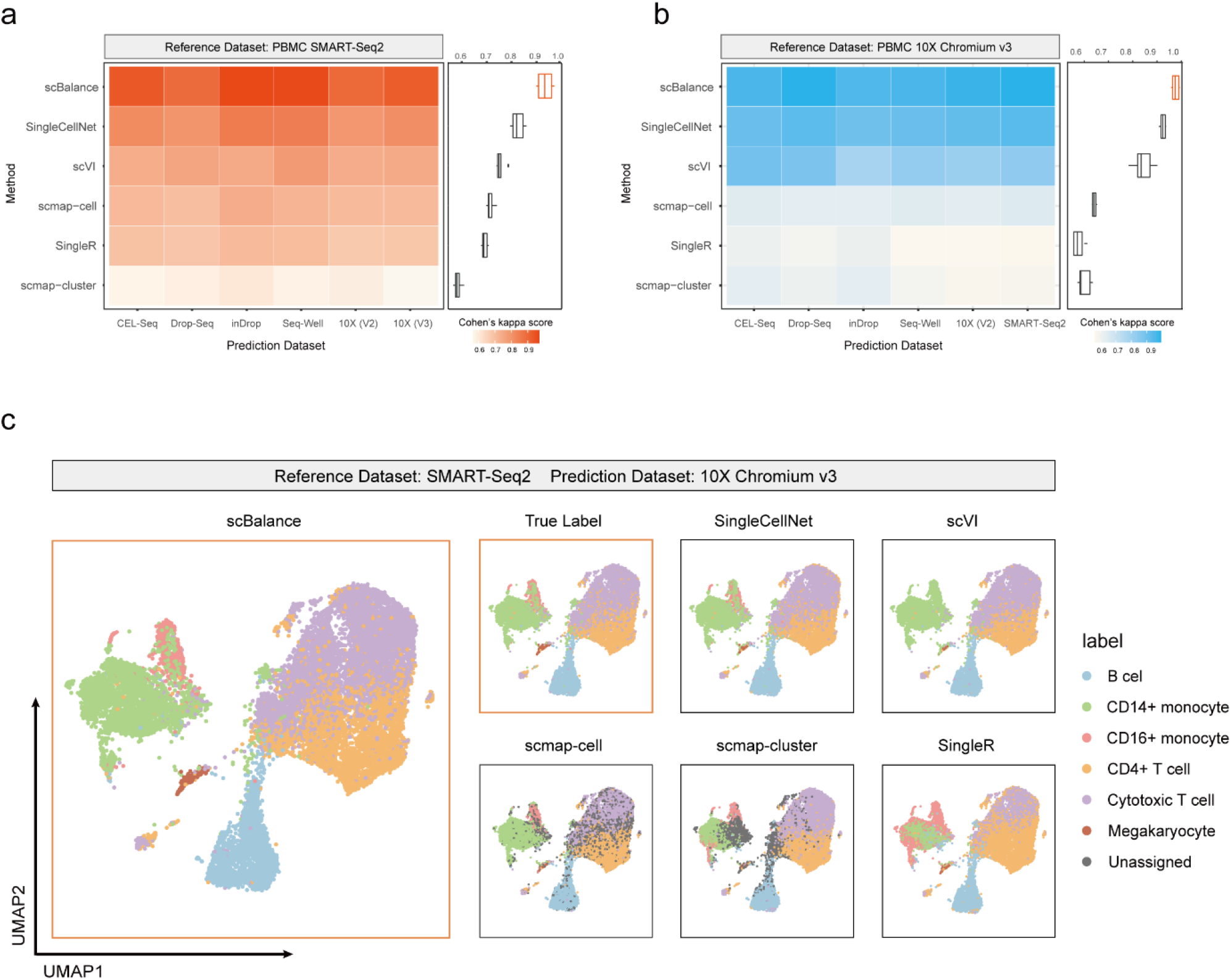
scBalance accurately identify rare cell types on the cross-platform annotation task. a. Overall annotation accuracy on the datasets generated by different protocols. scBalance outperforms all other five tools on the classification accuracy when with trained dataset generated from an alternative protocol to that used for the test. The performance is elevated by 10% than the second-best method. b. UMAP visualization showing the ability of identifying rare cell populations of different approaches. All methods are trained with the PBMC dataset (SMART-Seq2) and used to predict cell types in the PBMC dataset (10xv3). scBalance shows the best result identifying Megakaryocyte cells and CD16+ monocytes.

Meanwhile, because of the batch effect, annotating rare cell type can be more challenging. We notice that scBalance may also perform better on the rare subtype identification on the cross-platform tasks. To gain more insights into classification results of the rare population, we use Uniform Manifold Approximation and Projection (UMAP) to visualize the clustering result with the prediction label or true label (Fig.3b). Notably, we find that scBalance has the highest accuracy classifying the rare cell type. Compared with the true label, SingleCellNet gives more incorrect annotations on the Megakaryocyte cells and CD16+ monocytes than scBalance. Similarly, scVI also gives more incorrect labels on the Megakaryocyte cells and even completely fail on the classification of CD16+ monocytes. Though SingleR has a promising performance on labeling Megakaryocyte, it fails annotating CD14+ and CD16+ monocytes and has more mistakes on classifying Cytotoxic T cells and CD4+ T cells. In contrast, scBalance gives the most accurate annotation result on all six cell types and successfully labels the two rare cell populations, Megakaryocytes and CD16+ monocytes. Taken together, scBalance has a more robust performance on the cross-platform annotation tasks than the existing methods, and still has an outstanding capability on identifying rare cell population under the batch effect influence.

### Fast and robustness on the running speed enhances scalability of scBalance

Running time is considered as one of the most essential things for an annotation tool in the real single-cell analysis environment as well as the greatest obstacle of scalability. To highlight the superior of the scBalance on the calculation speed, we here present the comparison results of the running time of all the six methods (Fig. 4 and Supplementary Table 3). Because of the usage of the GPU, we separately show the scBalance-CPU and scBalance-GPU in order to make the comparison fair for other methods without GPU computation. We first compared the performance of the scBalance on the different processing units. The result indicates that scBalance-GPU has a large improvement on the running speed, which reduces more than 50% running time compared to the scBalance-CPU (Fig. 4a). Especially, scBalance-GPU gives a robust performance on the datasets with different cell numbers. The running time keeps relatively stable on the samples from 30k cells to 60k cells. This robustness gives scBalance a potential expanding ability to annotate large-scale datasets in a fast manner. We also present the comparison result of scBalance-CPU with the other five methods. Even all the methods are based on the CPU, scBalance also gives a promising running speed. Notably, in the datasets with more than 30k cells, scBalance reduced the running time to 10% of the other 5 methods. In the largest dataset, scBalance gives more than 20 times computation speed compared with SingleR (Fig. 4b). The advantage in time consuming also makes scBalance an outstanding tool in large-scale dataset annotation.

**Fig. 4.**
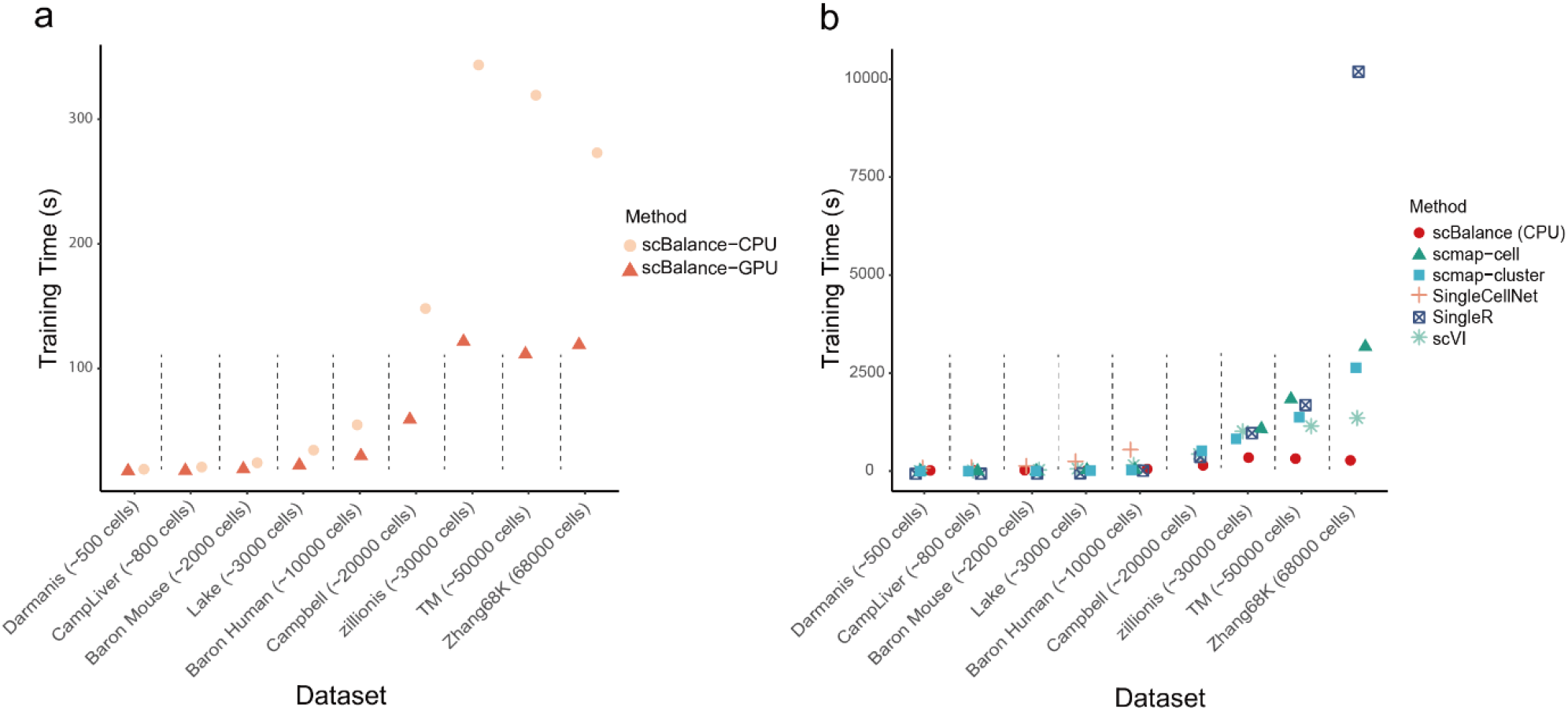
scBalance has the fastest running speed among these six methods. a. Comparison of the running time of scBalance on datasets of different scales. Different processers are used to run scBalance. b. Comparison of the running times of all the six methods on datasets of different scales. All methods are tested on the CPU.

### Revealing bronchoalveolar immune cell atlas in COVID patient proofs the scalability of scBalance

With the enlarging scale of the cell atlas, the scalability of an annotation tool becomes more and more important. We thus discuss the strength of scBalance to learn rare cell types in the million-level scRNA-seq datasets. We first use the intra-dataset annotation result as proof of concept to evaluate the annotation performance of scBalance on large scale cell atlas. We collected two recently published cell atlas including human heart cell atlas^31^ (487,106 cells) and COVID-19 immune atlas^17^ (1,462,702 cells). Because no other existing methods reported annotation ability on million-level scRNA-seq profiles, especially it is even hard to load the dataset for R-based methods such as SingleCellNet and Scmap, we compared scBalacne with conventional machine learning methods such as random forest (n_estimators=50,random_state=10), decision tree, SVM (kernel:rbf), and kNN (k=3) in Python. As shown in the Fig. 5a, scBalance significantly outperforms the other machine learning methods across both two cell atlases. Meanwhile, compared with the other methods, scBalance increases running speed up to at most 150 times when training and labeling COVID cell atlas (Fig. 5b). Even though the cell number increases 3 times between the two datasets, scBalance is the only method that can remain a robust running speed, which gives it more advantage in scalability (Supplementary Table 4-5).

**Fig. 5.**
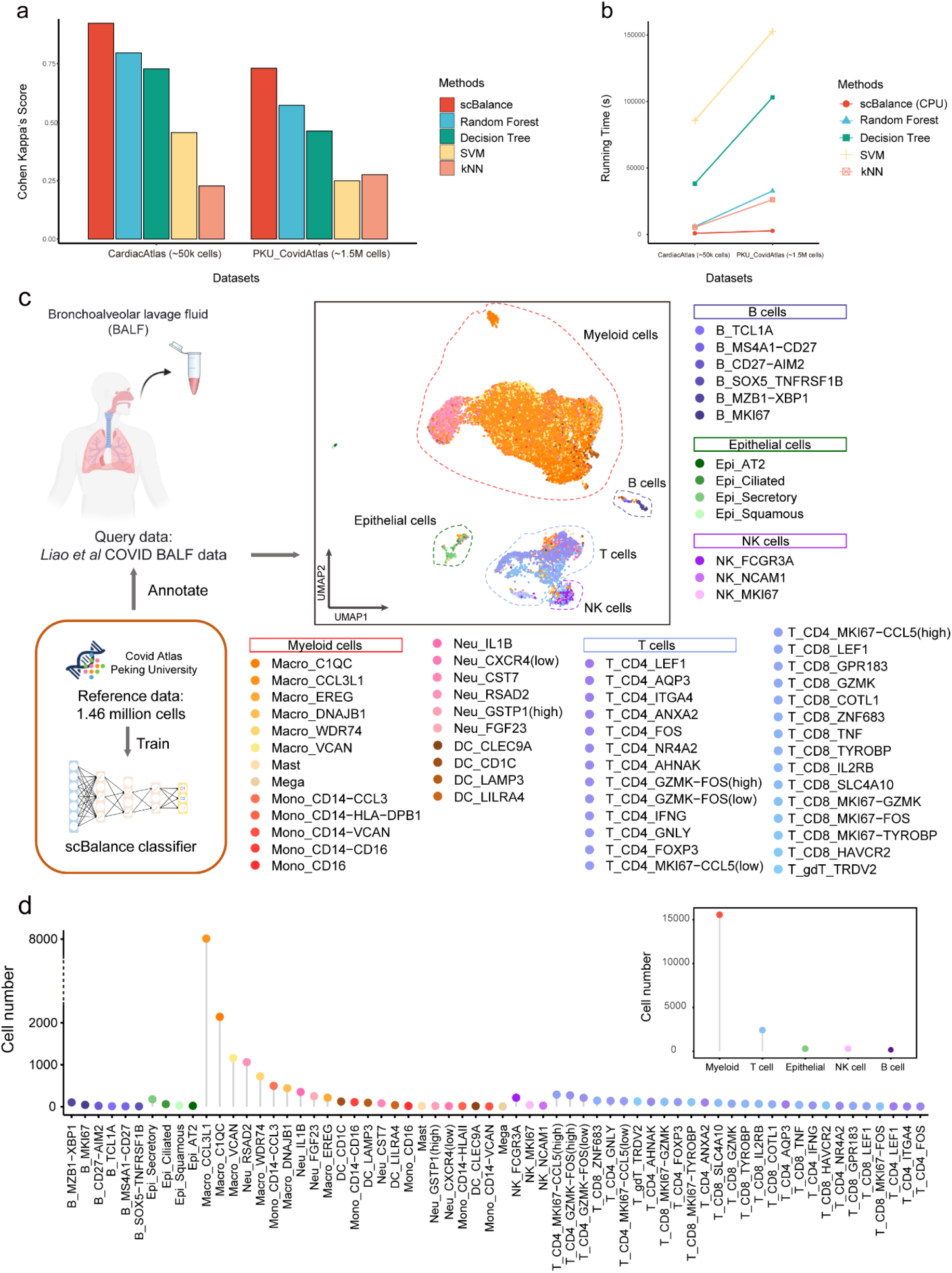
scBalance shows scalability by revealing immune landscape of BALF cells. **a**. annotation performances comparing with different methods on Cardiac Atlas (∼50K cells) and COVID Atlas (∼1.5M cells). **b**. Running time comparison between scBalance and traditional machine learning algorithms. Y-axis shows running time in second. **c**. UMAP shows the annotation result of scBalance. Reference dataset is COVID Atlas^17^ and query datatset is BALF data^28^. **d**. Dotplot shows the cell subtype distribution in the BALF dataset.

In addition to the simple evaluation of the scalability, we use COVID immune atlas as the reference dataset for an instance to illustrate that the annotation result of scBalance can effectively identify rare cell types when training with million scale references. We also collect Bronchoalveolar lavage fluid (BALF) cells scRNA-seq profile from a severe COVID patient as the query data (Fig. 5c). While there are lots of publications discussed PBMC landscape^32-35^ in different COVID patient samples, the BALF cell component of COVID patient is still lack investigation. But as the sample that can most directly reflect microenvironment information on lung alveoli, BALF cells are of great importance to understand the association of the disease severity and respiratory immune characteristics dynamic. Although *Liao et al* revealed bronchoalveolar immune cells landscape in patients with COVID in 2020^28^, their work which is based on the integration of seurat only identified cell groups in a low resolution. Here, we use scBalance to annotate BALF scRNA-seq dataset. Our method successfully identified much more cell subtypes in the datasets and yield a comprehensive immune landscape of BALF in the severe COVID patient. Compared to the original analysis, scBalance significantly improve annotation resolution for the BALF dataset. In combination with the result in the Fig. 5c, d and Supplementary Fig. 1, scBalance identified 64 subtypes of the immune cells in the BALF sample. As expected, macrophages show the highest enrichment in the BALF sample whereas B cells only be a small part of the immune landscape. Notably, scBalance also identified rare subtypes in all cell groups. In the myeloid group, our method elucidates that there are also monocyte locates in the BALF instead of only macrophage. But macrophage cells are still the major component, especially the proinflammatory macrophage (M1) such as CCL3L1^+^ macrophage, which suggests a strong immune cell recruitment signal in BALF in the severe patient. Meanwhile, different with the analysis by *Liao et al*^*28*^, our method reveals that the proinflammatory environment is not only produced by macrophage but also by CD14 monocyte (CCL3^+^). Furthermore, our method also finds that a significant expansion of proliferative memory T cells (including MKI67-CCL4 (high) CD4 T cell and MKI67-CCL4 (low) CD4 T cell), compared with effector T cells, are enriched in the lung region. Together, our methods successfully identify cell subtypes and provide a more comprehensive immune atlas in the BALF by using the COVID cell atlas as the reference. It is worth noting that most of the cell types revealed by scBalance are rare in the COVID atlas, which further presents the advantage of identifying rare cell types of our method in the large-scale scRNA-seq dataset.

## Discussion

Recent developments of scRNA-seq methods have increased the demand for cell-type annotation tools. With more and more well-defined cell atlases are being published, auto-annotation tool is becoming predominant. However, rare cell type labeling, scalability, and compatibility make limitations of the current software. In this article we present scBalance, an open-source Python package that integrate an adaptive weight sampling and a sparse neural network for supervised cell type auto-annotation. With intra- and inter-dataset comparison experiments on several scRNA-seq datasets with different scales, different generation protocols, and different imbalance degrees, we have shown an effective rare type annotation ability as well as a superior overall cell annotation ability of scBalance. We have also demonstrated a robust running speed of scBalance on different dimension datasets, which gives scBalance a potential advantage for scalability. In addition, by testing our method on two recently published large cell atlas, we have further shown the scalability and rare population identification capacity in million scale dataset of scBalance. By utilizing this ability, scBalance has successfully described an immune landscape of BALF cells and identified more rare types than the published research. Moreover, scBalance is specifically designed to be compatible with Scanpy and Anndata thus providing a user-friendly application.

We also suggest multiple future efforts for improving scBalance. Firstly, more information could be included as prior knowledge, such as marker genes, to make more accurate annotations for similar cells. Secondly, scBalance could be expanded to annotate single-cell chromatin accessibility sequencing (scATAC-seq) data by modifying network to a sparse-robust structure. Taken together, we believe that scBalance is an effective and important addition to the auto-annotation toolbox, especially by its rare cell type annotation ability and scalability.

## Methods

### Datasets

In this section, we describe all the datasets we used in the experiments and analysis above. In the baseline annotation experiment, we used 20 datasets from small scale (∼10k cells) to large scale (∼1.4M cells). To further demonstrate the generalization ability of scBalance, we select datasets with different complexities and different sequencing protocols. All the datasets and their corresponding cell type label are obtained from the original paper. The details are shown in Table 1.

### scBalance pipeline

We provide scBalance, a compounded neural network structure, to conduct cell type annotation tasks. scBalance requires a single-cell RNA expression matrix *M* as an input, in which each column represents a gene, and each row represents a cell. To obtain a more accurate annotation result, we recommend using a filtered dataset with log transformation and normalization as the training set. The goal is to prevent the outlier genes interfering training process. Preprocessing can be done by following the tutorial of Scanpy^8^, in which the scale parameter can be manually changed in the normalization function. The prediction dataset should have the same preprocessing steps as the training set. Before training, subsets will be extracted from the reference set and predicting set based on the common genes and be used as the input. scBalance pipeline consist of two core modules (Fig.1a), weighted sampling function and neural network classifier.

### Weighted sampling function

The first module is a weighted sampling function that provides a simple but efficient solution for the learning imbalanced scRNA-seq datasets. Unlike commonly used over-sampling and under-sampling methods, scBalance offers a combination of these two methods, thus significantly improving running speed without overfitting the minor types. In the training step, because we have the known labels in the training set, scBalance gives a weight to each cell type according to the proportion and randomly chooses samples from the dataset based on the weights to construct the training batch for the neural network. The sampling process is set with replacement to ensure the classifier can learn as much as possible minor type information in a reliable way.

### Neural network classifier

In the second module, we used a neural network (NN) structure to conduct the classification task. The NN classifier in scBalance contains an input layer, three hidden layers, and a softmax layer. The number of neurons in the input layer equals the number of genes in the scRNA-seq dataset. Following the three hidden layers have 256, 128, 64 units, respectively. We also add dropout and batch normalization techniques at each hidden layer to overcome overfitting and increase running speed. Only the training stage of scBalance involves forward propagation with Batch Normalization and Dropout techniques. To avoid the variance shift^36^, we put Dropout layer after Batch Normalization layer:

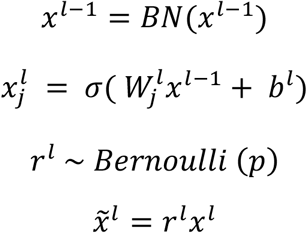

where *l* represents the *l*^*th*^ layer of the neural network, *j* represents the *j*^*th*^ neuron in its layer, b represents the random bias added in the layer, and *σ*(·) represents activation function. *BN*(·) is the batch normalization function to normalize the value of each mini-batch. *r* is a vector of independent Bernoulli random variable with the dropout probability *p*. This vector multiplied element-wise with each hidden layer to create dropout layer 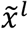. In scBalance, the default dropout probability is 0.5. The activation function in scBalance is exponential linear unit (ELU) function,

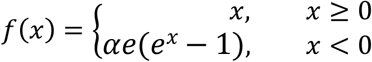

The output layer is based on the softmax function:

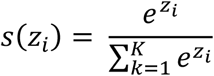

where *z* is the input vector of the softmax layer, *K* is the number of cell types in the reference dataset. In the backpropagation, we choose cross-entropy loss as the loss function of scBalance and the Adam^37^ optimization method as the optimizer. After training, the dropout layer will be disabled. scBalance provides a three-layer fully connected neural network for cell type prediction.

### Softeware comparison and settings

To testify the performance of scBalance, we compared it with several commonly used methods including R-based packages such as Scmap-cell, Scmap-cluster, SingleCellNet, SingleR, and Python-based package scVI. All the evaluation codes and input data follow the instructions and tutorials provided by each package. To ensure our evaluation is fair to each method, we set all parameters as default for each approach including scBalance.

The running environment we used for Python-based software is: (1) scVI from Github (https://github.com/YosefLab/scvi-tools). And the version is 0.14.5. We ran the GPU version and set the hyper-parameters following their example. We included LTMG inferring in preprocessing with the corresponding given option of the code. All the hyper-parameters are set following the tutorial. The task is implemented on the workstation with Intel(R) Xeon(R) CPU E5-2667 v4, CentOS Linux release 7.7.1908 operation system, Nvidia TITAN X GPU, 503GB physical memory. For the R-based packages, we implemented the tasks with the computer model Intel(R) Core(TM) i5-5287U CPU @ 2.90GHz RAM 8GB. The details of the softwares are: (2) SingleR version 1.6.1 from CRAN (https://github.com/dviraran/SingleR). The parameters are set as the default value provided by the tutorial. (3) SingleCellNet version 0.1.1 from BiocManager (https://github.com/pcahan1/singleCellNet), running with the default parameters. We took the category with the largest score in the prediction as the final result. It is worth mentioning that SingleCellNet can only deal with relative smaller datasets, and for the larger ones, it will stuck while running. (4) Scmap-Cell and Scmap-Cluster from BioManager (https://github.com/hemberg-lab/scmap), with all parameters following the function instruction.

### Performance evaluation

We describe below the quantitative metrics we used in the experiments. To evaluate the classification accuracy of the scBalance, we used Cohen’s kappa score and F1 score. Unlike most of the papers which use Accuarcy (Acc) as the metric, our aim is to testify the identification ability of the rare cell types as well as the overall classification accuracy. Therefore, we choose Cohen’s kappa coefficient^38^ *k*, which is a minor-class sensitive approach thus can give us a comprehensive evaluation of classification performance including the major types identification and the minor types indentifcation,

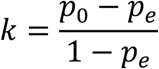

where *p*_0_ is the observed proportionate variable and *p*_*e*_ is the hypothetical probability of chance variable. To calculate *p*_*e*_, we use the observed data to calculate the probabilities of each observer randomly seeing each category. In this formula, the weight for misclassification of the rare populations will be highlighted.

## Data availability

No new data was generated for this study. All data used in this study is publicly available as previously described (see Table 1).

## Code availability

scBalance is available as an independent python package at https://github.com/yuqcheng/scBalance.

## Contributions

Y.C. designed the method and all benchmark experiments. Y.C. implemented the tool in Python. Y.C., J.Z., and X.F. performed data analysis and all computation experiments. J.Z. and X.F. also provided advice in method development. Y.C., Y.L.,J.Z., and X.F. wrote the manuscript together. All author reviewed the manuscript.

## Competing interests

The authors declare no competing interests.

---

**Supplementary Fig.1.**
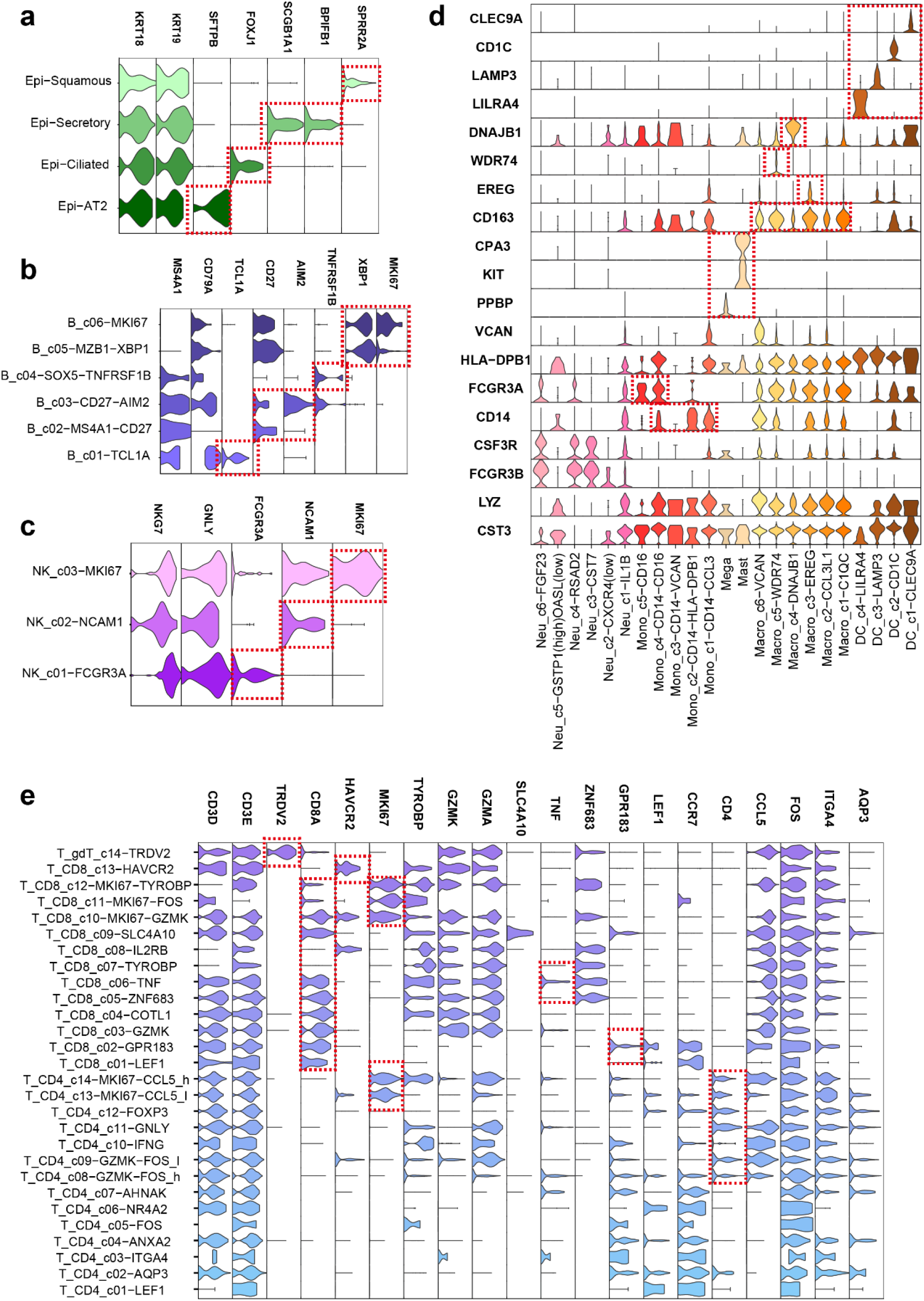
Violin plots show the selected marker genes for each identified cell type. Including **a**. 4 epithelial cell types, **b**. 6 B cell types, **c**. 3 NK cell types, **d**. 23 Myeloid cell types and **e**. 28 T cell types.

## Notes

### Competing Interest Statement

The authors have declared no competing interest.

## Reference

1. Tang, F. et al. mRNA-Seq whole-transcriptome analysis of a single cell. Nature Methods 6, 377–382 (2009).

2. Horning, A.M. et al. Single-Cell RNA-seq Reveals a Subpopulation of Prostate Cancer Cells with Enhanced Cell-Cycle–Related Transcription and Attenuated Androgen Response. Cancer Research 78, 853–864 (2018).

3. Nyquist, M.D. et al. Combined TP53 and RB1 Loss Promotes Prostate Cancer Resistance to a Spectrum of Therapeutics and Confers Vulnerability to Replication Stress. Cell Reports 31, 107669 (2020).

4. Guo, C. et al. Single-cell analysis of two severe COVID-19 patients reveals a monocyte-associated and tocilizumab-responding cytokine storm. Nature Communications 11 (2020).

5. Wilk, A.J. et al. A single-cell atlas of the peripheral immune response in patients with severe COVID-19. Nature Medicine 26, 1070–1076 (2020).

6. Guo, L. et al. Resolving Cell Fate Decisions during Somatic Cell Reprogramming by Single-Cell RNA-Seq. Molecular Cell 73, 815–829.e817 (2019).

7. Butler, A., Hoffman, P., Smibert, P., Papalexi, E. & Satija, R. Integrating single-cell transcriptomic data across different conditions, technologies, and species. Nature Biotechnology 36, 411–420 (2018).

8. Wolf, F.A., Angerer, P. & Theis, F.J. SCANPY: large-scale single-cell gene expression data analysis. Genome Biology 19 (2018).

9. Tran, D. et al. Fast and precise single-cell data analysis using a hierarchical autoencoder. Nature Communications 12 (2021).

10. Han, W. et al. Self-supervised contrastive learning for integrative single cell RNA-seq data analysis. BioRxiv (2021).

11. Lindeboom, R.G.H., Regev, A. & Teichmann, S.A. Towards a Human Cell Atlas: Taking Notes from the Past. Trends in Genetics 37, 625–630 (2021).

12. Rozenblatt-Rosen, O., Michael, J., Regev, A. & Teichmann, S.A. The Human Cell Atlas: from vision to reality. Nature 550, 451–453 (2017).

13. Xie, B., Jiang, Q., Mora, A. & Li, X. Automatic cell type identification methods for single-cell RNA sequencing. Computational and Structural Biotechnology Journal 19, 5874–5887 (2021).

14. Tan, Y. & Cahan, P. SingleCellNet: A Computational Tool to Classify Single Cell RNA-Seq Data Across Platforms and Across Species. Cell Systems 9, 207–213.e202 (2019).

15. Ma, F. & Pellegrini, M. ACTINN: automated identification of cell types in single cell RNA sequencing. Bioinformatics (2019).

16. Worbs, T., Hammerschmidt, S.I. & Förster, R. Dendritic cell migration in health and disease. Nature Reviews Immunology 17, 30–48 (2017).

17. Ren, X. et al. COVID-19 immune features revealed by a large-scale single-cell transcriptome atlas. Cell 184, 5838 (2021).

18. Khalilia, M., Chakraborty, S. & Popescu, M. Predicting disease risks from highly imbalanced data using random forest. BMC Medical Informatics and Decision Making 11, 51 (2011).

19. Zheng, G.X.Y. et al. Massively parallel digital transcriptional profiling of single cells. Nature Communications 8, 14049 (2017).

20. Han, X. et al. Mapping the Mouse Cell Atlas by Microwell-Seq. Cell 172, 1091–1107.e1017 (2018).

21. Nguyen, V. & Griss, J. scAnnotatR: framework to accurately classify cell types in single-cell RNA-sequencing data. BMC Bioinformatics 23 (2022).

22. Zhang, R., Luo, Y., Ma, J., Zhang, M. & Wang, S. scPretrain: Multi-task self-supervised learning for cell type classification. BioRxiv (2020).

23. Wang, L. et al. An interpretable deep-learning architecture of capsule networks for identifying cell-type gene expression programs from single-cell RNA-sequencing data. Nature Machine Intelligence 2, 693–703 (2020).

24. Kim, H., Lee, J., Kang, K. & Yoon, S. (Research Square Platform LLC, 2021).

25. Kiselev, V.Y., Yiu, A. & Hemberg, M. scmap: projection of single-cell RNA-seq data across data sets. Nature Methods 15, 359–362 (2018).

26. Aran, D. et al. Reference-based analysis of lung single-cell sequencing reveals a transitional profibrotic macrophage. Nature Immunology 20, 163–172 (2019).

27. Lopez, R., Regier, J., Cole, M.B., Jordan, M.I. & Yosef, N. Deep generative modeling for single-cell transcriptomics. Nature Methods 15, 1053–1058 (2018).

28. Liao, M. et al. Single-cell landscape of bronchoalveolar immune cells in patients with COVID-19. Nature Medicine 26, 842–844 (2020).

29. Srivastava, N., Hinton, G., Krizhevsky, A., Sutskever, I. & Salakhutdinov, R. Dropout: a simple way to prevent neural networks from overfitting. The journal of machine learning research 15, 1929–1958 (2014).

30. Clevert, D.-A.e., Unterthiner, T. & Hochreiter, S. Fast and Accurate Deep Network Learning by Exponential Linear Units (ELUs). arXiv (2016).

31. Litvinukova, M. et al. Cells of the adult human heart. Nature 588, 466-+ (2020).

32. Wilk, A.J. et al. A single-cell atlas of the peripheral immune response in patients with severe COVID-19. Nat Med 26, 1070–1076 (2020).

33. Schulte-Schrepping, J. et al. Suppressive myeloid cells are a hallmark of severe COVID-19. medRxiv, 2020.2006.2003.20119818 (2020).

34. Zhao, J. et al. Antibody Responses to SARS-CoV-2 in Patients With Novel Coronavirus Disease 2019. Clinical Infectious Diseases 71, 2027–2034 (2020).

35. Rabaan, A.A. et al. Role of Inflammatory Cytokines in COVID-19 Patients: A Review on Molecular Mechanisms, Immune Functions, Immunopathology and Immunomodulatory Drugs to Counter Cytokine Storm. Vaccines (Basel) 9 (2021).

36. Li, X., Chen, S., Hu, X. & Yang, J. Understanding the Disharmony between Dropout and Batch Normalization by Variance Shift. arXiv (2018).

37. Diederik & Ba, J. Adam: A Method for Stochastic Optimization. arXiv (2017).

38. Vieira, S.M., Kaymak, U. & Sousa, J.M.C. Cohen’s kappa coefficient as a performance measure for feature selection. International Conference on Fuzzy Systems (2005).

39. Deng, Q.L., Ramskold, D., Reinius, B. & Sandberg, R. Single-Cell RNA-Seq Reveals Dynamic, Random Monoallelic Gene Expression in Mammalian Cells. Science 343, 193–196 (2014).

40. Darmanis, S. et al. A survey of human brain transcriptome diversity at the single cell level. Proceedings of the National Academy of Sciences 112, 7285–7290 (2015).

41. Usoskin, D. et al. Unbiased classification of sensory neuron types by large-scale single-cell RNA sequencing. Nature Neuroscience 18, 145–153 (2015).

42. Camp, J.G. et al. Multilineage communication regulates human liver bud development from pluripotency. Nature 546, 533–538 (2017).

43. Baron, M. et al. A Single-Cell Transcriptomic Map of the Human and Mouse Pancreas Reveals Inter-and Intra-cell Population Structure. Cell Systems 3, 346–360.e344 (2016).

44. Mauro et al. A Single-Cell Transcriptome Atlas of the Human Pancreas. Cell Systems 3, 385–394.e383 (2016).

45. Lake, B.B. et al. Neuronal subtypes and diversity revealed by single-nucleus RNA sequencing of the human brain. Science 352, 1586–1590 (2016).

46. Campbell, J.N. et al. A molecular census of arcuate hypothalamus and median eminence cell types. Nature Neuroscience 20, 484–496 (2017).

47. Zilionis, R. et al. Single-Cell Transcriptomics of Human and Mouse Lung Cancers Reveals Conserved Myeloid Populations across Individuals and Species. Immunity 50, 1317-+ (2019).

48. Schaum, N. et al. Single-cell transcriptomics of 20 mouse organs creates a Tabula Muris. Nature 562, 367-+ (2018).

49. Zheng, G.X.Y. et al. Massively parallel digital transcriptional profiling of single cells. Nature Communications 8 (2017).

50. Ding, J. et al. Systematic comparison of single-cell and single-nucleus RNA-sequencing methods. Nature Biotechnology 38, 737–746 (2020).

